# Molecular noise accelerates cell division and death under antibiotic treatment

**DOI:** 10.1101/124552

**Authors:** Savita Chib, Suman Das, V. Soumya, Aswin Sai Narain Seshasayee, Mukund Thattai

## Abstract

It is well known that microbial cell populations can exhibit sustained exponential growth. More surprising is the fact that at high antibiotic levels, cell populations exhibit sustained exponential decay over several orders of magnitude. The boundary between growth and decay occurs at the Minimal Inhibitory Concentration (MIC) of the antibiotic, where the density of living cells remains constant over time. These observations suggest that positive (growth) or negative (decay) exponents arise as a difference of cell division and death rates obeying first-order kinetics. Thus for antibiotic concentrations below MIC, division dominates; for concentrations above MIC, death dominates; while MIC itself is a dynamic steady state of balanced division and death, rather than cell stasis. To measure these rates we separately tracked living and dead cells in *Escherichia coli* populations treated with the ribosome-targeting antibiotic kanamycin. We found that cells divide rapidly even at MIC: inferred division and death rates at MIC are 0.6 times the antibiotic-free division rate. A stochastic model of cells as collections of self-replicating units we term “widgets” reproduces both steady-state and transient features of our experiments, and explains first-order exponential kinetics. In this model cell division and death rates at MIC can be tuned from low to high values by amplifying molecular noise in the synthesis, degradation and partitioning of the widgets. At extremely low noise, cells approach the classic bacteriostatic limit at MIC: neither dividing nor dying. Noise-induced division and death of cells following antibiotic treatment could increase the likelihood of sepsis and antibiotic resistance.

---

Exponential growth and decay are macroscopically observable features of cell populations; division and death are microscopic single-cell processes. It is tacitly assumed that growing populations contain mainly dividing cells, while decaying populations contain mainly dying cells. The reality is richer and more subtle: under many growth conditions, cells are both dividing and dying, and the net growth or decay exponent arises from the difference of the two [1: Liu et al., 2004]. This idea forms the starting point of our discussion, and naturally leads us to confront randomness and molecular noise.

The study of bacterial growth has been instrumental in revealing fundamental aspects of cell division, metabolism, and regulation [2: Monod, 1949; 3: Campbell, 1957; 4: Zaritsky & Woldringh, 2015]. Campbell [3: 1957] realized that exponentially growing populations were in a state of “balanced growth”: the chemical composition of a daughter cell immediately after division was invariant from one generation to the next, leading to a well-defined and constant doubling time. The specific dependence of the exponential growth rate (the exponent of the cell density versus time curve) on nutrient or antibiotic concentrations can be summarized as “growth laws” [5: Schaechter et al., 1958]. Models of bacteria as autocatalytic chemical reactors accurately capture many mathematical features of these growth laws [6: Scott et al., 2010; 7: Scott et al., 2014; 8: Parth & Jain, 2017].

Studies of bacterial growth laws have focused mainly on exponential growth. However, bacterial populations in the presence of high antibiotic levels can also undergo sustained exponential decay over several orders of magnitude (Fig. 1A) [9: Craig & Ebert, 1990; 10: Barbour et al., 2010]. This is surprising: exponential growth can arise from deterministic cell doubling, but exponential decay with first-order kinetics typically occurs when individuals in a population die at random, like radioactive nuclei. However, cells are not indivisible objects: if some cells die early while others die later, this must be due to some underlying cell-to-cell variability. The growth of single cells is known to be a stochastic, fluctuating process [11: Kiviet et al., 2014; 12: Iyer-Biswas et al., 2014; 13: Taheri-Araghi et al., 2015], an outcome of noisy biochemical reactions [14: Raj & van Oudenaarden, 2008]. We should therefore consider the possibility that the exponential growth or decay of a cell population arises as a competition between two random microscopic processes: cell division and cell death. This contrasts with the usual assumption that any change in exponential growth rates is due to a change in division rate alone, rather than a difference of division and death rates [6: Scott et al., 2010; 7: Scott et al., 2014]. No analysis has so far attempted to simultaneously account for molecular noise, cell division, and cell death within a common framework.

**Figure 1:**
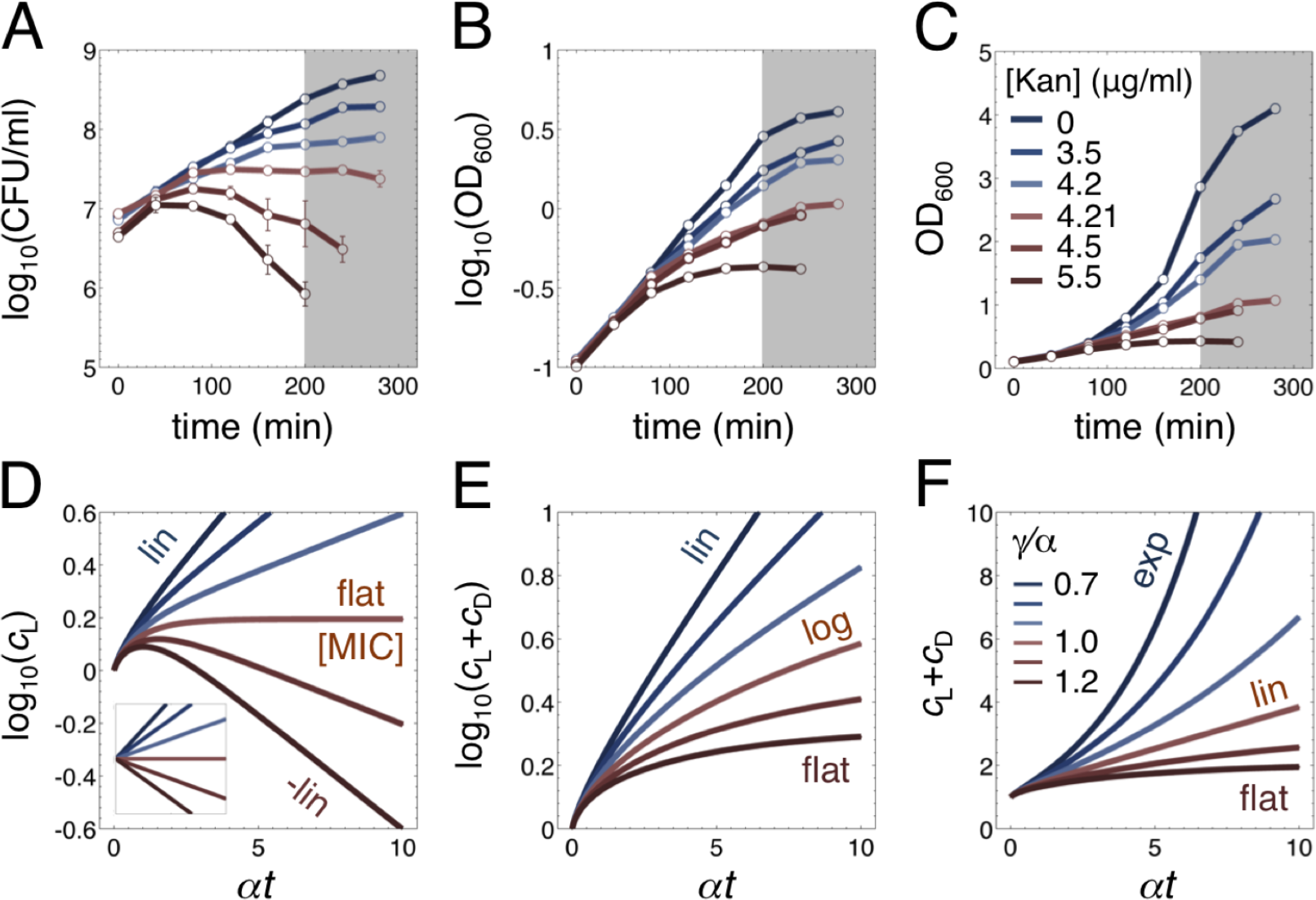
Growth and decay of cell populations under antibiotic treatment. (**A,B,C**) Measurements for *E. coli* populations. Each curve corresponds to different concentrations of the antibiotic kanamycin; see key in panel (C). [Kan] = 4.21 μg/ml is the Minimal Inhibitory Concentration (MIC) at which the viable cell count is constant over time. (**A**) Viable cell count (CFU/ml) over time, mean and standard deviation over four technical replicates. (**B,C**) Turbidity (OD_600_) over time, on logarithmic and linear axes. The gray area indicates the nutrient depletion zone. (**D,E,F**) Predictions of a stochastic model of cell growth. Curves show solutions to Eq. 4 for the division threshold *Ω* = 10 and increasing values of *γ*/*α*, corresponding to increasing antibiotic levels; see key in panel (F). *γ*/*α* = 1 corresponds to MIC. (**D**) The number of living cells *c_L_* corresponds to CFU/ml. The inset shows the prediction for a cell that has no internal structure and divides as soon as *Ω* = 2. (**E,F**) The total number of living and dead cells *c_L_* + *c_D_* corresponds to OD_600_. We have labeled the asympotic behavior of the predicted curves: exponential (exp), linear (lin), logarithmic (log), or flat.

Here we experimentally demonstrate that growth at “bacteriostatic” antibiotic concentrations, where the total number of living cells remains constant over time, is in fact a dynamic balance between cell division and cell death. This observation can be understood if we think of cells as collections of sub-cellular self-replicating units. A self-replicating unit is a minimal autocatalytic set of molecules and reactions [15: Jain & Krishna, 2001]. The ribosome lies at the heart of such a unit, since most of the energy and biomass budget of a bacterial cell is devoted to ribosomes making more ribosomes [6: Scott et al., 2010; 7: Scott et al., 2014]. Ribosomes must be supplemented by DNA replication, and the synthesis of non-ribosomal proteins and biomass components via central metabolism. Here we sidestep the precise molecular nature of the self-replicating unit, and group it into a single abstract “widget”. We model the synthesis, degradation and partitioning of widgets as noisy biochemical processes. Cell division or death occurs when a cell hits high or low thresholds of these widgets. Remarkably, this basic model reproduces observed qualitative features of cell growth and decay, and gives a mechanistic explanation for the continued division of cells at high antibiotic concentrations. Our central insight is that molecular noise in the number of widgets can drive a stochastic choice between cell division and death. Increased levels of noise drive increased rates of both division and death, and the net number of living cells at any time is determined by the balance between these processes. Our observations raise fundamental questions about the molecular nature of the self-replicating widget, and of the blurry boundary between cell division and death.

## Results

### Measuring cell growth in the presence of antibiotics

Different classes of antibiotics act through distinct mechanisms [16: Kohanski et al., 2010]. Since we focus on self-replicating units, here we use the aminoglycoside antibiotic kanamycin which irreversibly binds to and inhibits the ribosome [17: Greulich et al., 2015]. The effect of an antibiotic is typically quantified in terms of its impact on growth rates. It is common to use turbidity measurements (OD_600_) for this purpose, since these are easy to carry out and automate [18: Toprak et al., 2013]. Cell growth is more accurately determined by measuring the density of viable colony-forming units (CFU/ml) [10: Barbour et al., 2010]. These two are not equivalent: colony-forming units measure the density of living cells, whereas turbidity measures the total density of all non-lysed cells, living or dead. The Minimal Inhibitory Concentration (MIC) of an antibiotic is often defined as the concentration at which turbidity increase is prevented, but this depends on the duration and sensitivity of the measurement [19: Chait et al., 2007]. Here we rigorously define MIC as the antibiotic concentration at which the density of colony-forming units is asymptotically constant over time. This “bacteriostatic” antibiotic treatment is sometimes contrasted with “bactericidal” treatments which cause cell death, but this is not a sharp distinction [16: Kohanski et al., 2010]. As we show below, at the microscopic level cell division and cell death can continue while the macroscopic count of viable cells remains static.

### Decomposing exponential growth or decay into cell division and death rates

We monitored the effect of kanamycin addition on *Escherichia coli* cells grown in an initially antibiotic-free medium (Fig. 1A,B,C; Methods: Growth and measurement protocols). To get a detailed picture of the effect of the antibiotic, we simultaneously measured living cell density (viability: CFU/ml) and total cell density (turbidity: OD_600_) over time. After a brief transient, CFU/ml settled into an exponentially growing profile (for low [Kan]) or an exponentially decaying profile (for high [Kan]) (Fig. 1A). At the boundary between these two regimes CFU/ml remained constant over time, defining the MIC ([Kan] = 4.21 μg/ml). The behavior of turbidity was strikingly different: OD_600_ always monotonically increased, with a convex accelerating profile (for low [Kan]) or a concave decelerating profile (for high [Kan]) (Fig. 1B,C). At the boundary between these two regimes, OD_600_ increased precisely linearly (Fig. 2B). These trends persisted until the medium was depleted of nutrients (gray zone, Fig. 1A,B,C).

**Figure 2:**
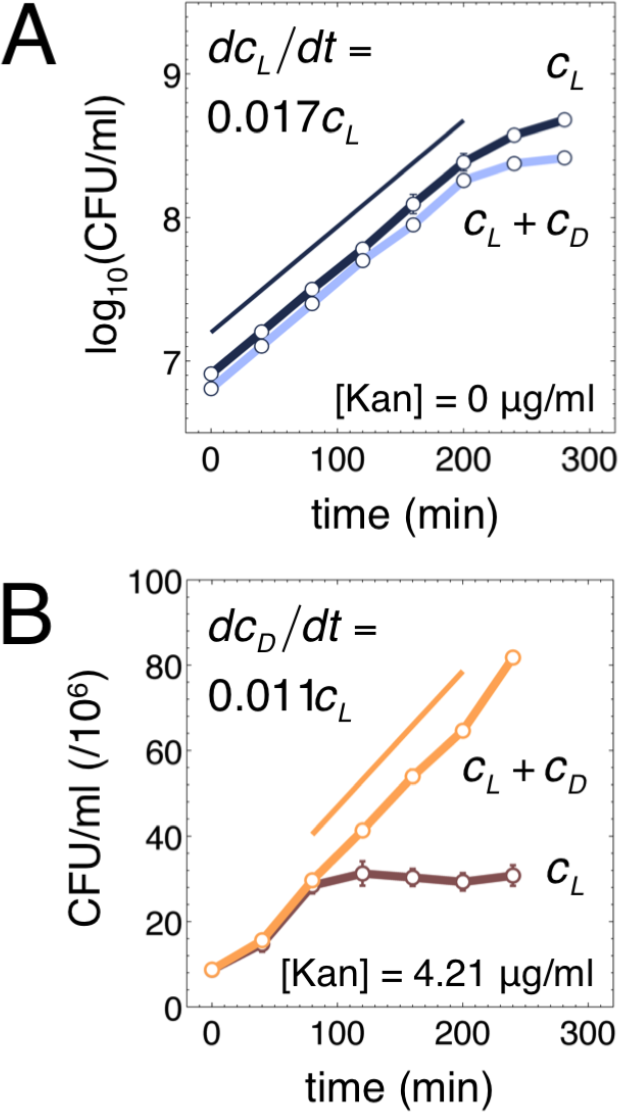
Comparing cell viability measurements (CFU/ml) to turbidity measurements (OD_600_) reveals cell dynamics at MIC. Dark curves: CFU/ml, equivalent to living cell number *c_L_* in Eq. 2. Light curves: OD_600_, equivalent to total cell number *c_L_* + *c_D_* in Eq. 2. We have applied the scale factor CFU/ml = 8 × 10^7^ × OD_600_ to plot both on the same axis. (**A**) At zero antibiotic CFU/ml and OD_600_ are completely overlapping and show perfect exponential growth in this log-lin plot; we have inserted an offset so both can be seen. The division rate is 0.017 min^−1^. (**B**) At MIC the CFU/ml curve flattens out while the OD_600_ curve continues to increase linearly in this lin-lin plot. Division and death rates are both equal to 0.011 min^−1^.

These observations are consistent with the idea that CFU/ml measures the density of living cells *c_L_*, which can increase or decrease) while OD_600_ measures the total density of living plus dead cells (*c_L_* + *c_D_*, which can only increase). As a first attempt to understand these dynamics, we decompose the separate contributions of cell division (*ϕ*_+_) and death (*ϕ*_−_) rates (Fig. 3C):

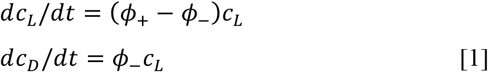
In the period once the transient response to the antibiotic has settled but nutrients are not yet depleted, *ϕ*_±_ are constant over time. The solution to Eq. 1 is then:

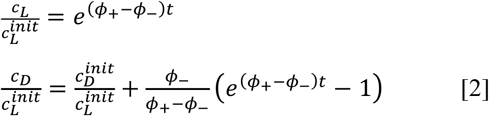
where we have scaled cell numbers by their initial values at the end of the transient.

**Figure 3:**
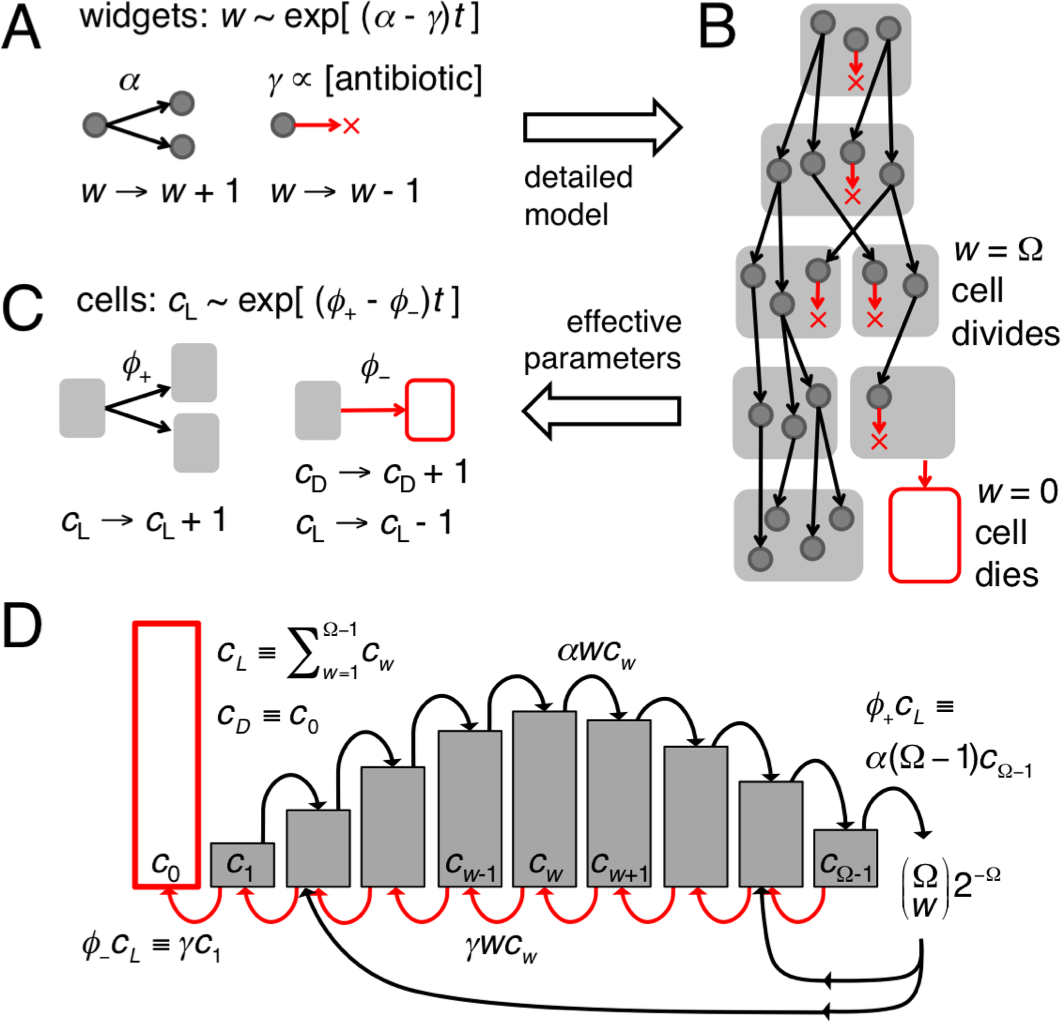
A stochastic model of cell division and death. (**A**) A widget is a self-replicating unit obeying a birth-death process with rates *α* and *γ*, the latter proportional to antibiotic levels. (**B**) Cells are collections of widgets. When a cell hits *w* = *Ω* it divides; when it hits *w* = 0 it dies. (**C**) The widget dynamics can be used to define the cell dynamics, with effective time-dependent division and death rates *ϕ*_±_. (**D**) Dynamics of a cell population: *c_w_* represents the number of cells with exactly *w* widgets. Individual cells move to the right (gain a widget) or left (lose a widget). *ϕ*_−_ is the per-cell rate at which cells cross the left boundary at *w* = 1 and die. *ϕ_+_* is the per-cell rate at which cells cross the right boundary at *w* = *Ω* − 1 and divided. At division the widgets binomially partition into two daughter cells, which re-enter the main distribution. The area of the red bin gives the number of dead cells, the total area of the gray bins gives the number of living cells. Over time the gray distribution reaches a constant shape but can increase or decrease in area.

*ϕ*_±_ depend on the antibiotic concentration, two values of which are of particular interest. At zero antibiotic we have 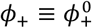 and *ϕ*_−_ = 0, so 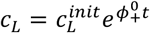 and *c_D_* = 0. Indeed we find OD_600_ is directly proportional to CFU/ml at zero antibiotic, and both grow with exponent 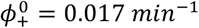 (or *τ_double_* ∼ 40 min; Fig. 2A). The scale factor 8 × 10^7^ converts OD_600_ to CFU/ml, allowing us to plot *c_L_* and *c_L_* + *c_D_* on the same axes. At MIC the situation is more interesting since by definition *ϕ*_+_ = *ϕ*_−_ = *ϕ*^MIC^, so 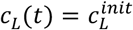 and 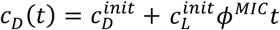. That is, living cell numbers are constant because death and division rates balance, while dead cell numbers increase linearly because they arise from the continuing death of living cells. This is precisely what we observe: the slope of the OD_600_ curve at MIC shows that *ϕ^MIC^* = 0.011 *min*^−1^ (or *τ_double_* ∼ 60 min; Fig. 2B). Interestingly the OD_600_ curve tracks the CFU/ml curve for the first hour following antibiotic treatment, after which the CFU/ml curve flattens while the OD_600_ curve increases linearly. This shows that cell death only begins after a lag, while cell division is relatively unperturbed by the presence of antibiotic. This is corroborated by the ratio 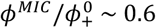 being close to unity: rapid cell division and death continue even at MIC.

### A model of cell growth in terms of widgets: sub-cellular self-replicating units

The initial transient response of cells to antibiotic addition is a sign of inertia due to internal re-organization. The phenomenological model of Eq. 1 incorrectly predicts pure exponential growth or decay (inset, Fig. 1D) because it assumes a cell has no internal structure. If we wish to determine how *ϕ*_±_ depend on antibiotic concentration or time, this must either be directly measured, or predicted from a more microscopic model. We therefore consider a cell as a collection of self-replicating units we term “widgets”. (Fig. 3; Methods: Stochastic model of cell division and death). The widgets themselves obey a birth-death dynamics analogous to Eq. 1, but with microscopic birth and death rate constants *α* and *γ* (Fig. 3A). For concreteness we imagine *α* to be constant (e.g. the catalytic efficiency of ribosomes) while *γ* depends on the antibiotic concentration (e.g. the rate of irreversible ribosome inhibition by kanamycin), but these assumptions may be relaxed. We specify how cell division and death depend on the widgets as follows (Fig. 3B). When the widget number hits *w* = 0 the cell dies since no new widgets can be made. When the widget number hits a threshold *w* = *Ω* the cell instantaneously divides, and the widgets are partitioned binomially between two daughter cells. This is arguably the simplest possible microscopic model of cell growth.

The dynamics of the widgets can be used to determine the effective parameters *ϕ*_±_ that appear in Eq. 1 (Fig. 3C). We consider a population of cells, binned according to the number of widgets they contain: *c_w_* is the number of cells with precisely *w* widgets, for *w* = 1,…, *Ω* − 1 (Fig. 3D; Methods: Stochastic model of cell division and death). Individual cells do a biased Poisson random walk along the *w*-axis: moving to the right if they gain a widget, or to the left if they lose one. The number of living and dead cells are:

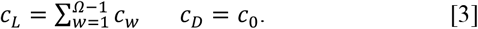

Cells that cross the left boundary (*i* = 1) move to bin *c*_0_ and die, so *c_L_* decreases by one. Cells that cross the right boundary (*i* = *Ω* − 1) move to the bin *c_Ω_* and instantaneously divide into two daughters, so *c_L_* increases by one. The daughters re-enter the distribution at two positions *w*′ and *w*″ such that *w*′ + *w*″ = *Ω*. These processes define a transition matrix *A* (Methods: Stochastic model of cell division and death) so that:

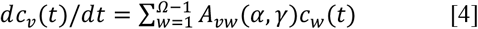

This can be easily solved for *c_w_*(*t*) from any *c_w_*(*t* = 0) by matrix exponentiation. Once we find the number of cells in each bin, we can track how many cells cross the right or left boundary. We can thus write an equation similar to Eq. 1, where *ϕ*_±_ are now time-dependent because the distribution of cells evolves from its initial state:

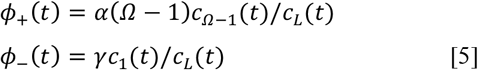

Over time, the distribution of living cells over widget number reaches a steady state, proportional to the eigenvector of the transition matrix corresponding to its largest eigenvalue *λ*. If we define a normalized distribution *f_w_*(*α, γ*), such that *A**f*** = *λ**f*** then:

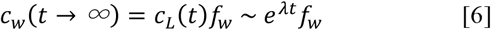

That is, the shape of the distribution becomes constant, while the total number of cells increases or decreases exponentially. In particular, this means that *ϕ*_±_ become time-independent, so comparing Eq. 6 to Eq. 2 we must have *λ* = *ϕ*_+_ − *ϕ*_−_. On the other hand, it is easy to check that the largest eigenvalue of *A* is given by *λ* = *α* − *γ* (Methods: Stochastic model of cell division and death). This gives us two completely distinct ways to decompose the growth exponent *λ* in the limit *t* → *∞*: in terms of widget birth/death, or in terms of cell division/death:

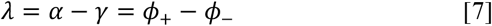

The intuition behind this result is simple: if the mean number of widgets per cell reaches a finite limit, cell number must be proportional to widget number. The interesting point is that the split between *α* and *γ* is very different than the split between *ϕ*_+_ and *ϕ*_−_.

### Comparison of widget model to experimental results

If we measure time in units of *α*^−1^ the model has only two dimensionless parameters: the widget death/birth ratio *γ*/*α*, and the threshold number of widgets at cell division *Ω*. Sweeping *γ*/*α* from low to high values corresponds to increasing the antibiotic concentration, with *γ*/*α* = 1 at MIC. The parameter *Ω* has a less direct interpretation, we will return to this when we discuss the role of molecular noise. In our experiments we first grow cells in in the absence of antibiotic and then add kanamycin at the initial measurement point. To model this we first find the cell distribution at zero antibiotic: *f_w_* (*α* = 1, *γ* =0). Starting with this *f_w_* as the initial condition, we use Eq. 4 to find *c_w_* (*t*) and Eq. 3 to find *c_L_*(*t*) and *c_D_*(*t*) for various ratios *γ*/*α*. The predictions using *Ω* = 10 (Fig. 1D,E,F) qualitatively match our experimental observations (Fig. 1A,B,C). In particular, we capture the initial transient bump as well as the asymptotic exponential growth and decay kinetics of CFU/ml (Fig. 1A,D). More interestingly, we also capture the response at MIC, where CFU/ml flattens while OD_600_ approaches a linear trajectory (Fig. 1B,C vs. Fig. 1E,F).

### Noise-induced cell division and cell death

The qualitative agreement of the model with measurements encouraged us to probe the underlying structure of the model further. The key goal is to understand what drives the choice of a post-division cell between two ultimate fates: division and death (Methods: Probability of division). Immediately following division the widgets partition binomially between daughter cells, leading to an initial post-division variation (histogram, Fig. 4A). At this point, the noisy birth-death widget dynamics take over. Cells with an initially low value of *w* are more likely to hit the left boundary and die, while cells with an initially high value are more likely to hit the right boundary and divide (curves, Fig. 4A). The addition of antibiotics (increasing *γ*/*α*) biases the choice against division. The squared coefficient of variation (SCV) of the post-division binomial distribution 1/*Ω* is a convenient measure of noise. Note that increasing 1/*Ω* increases the strength of molecular noise in both binomial widget partitioning and Poisson birth/death dynamics.

**Figure 4:**
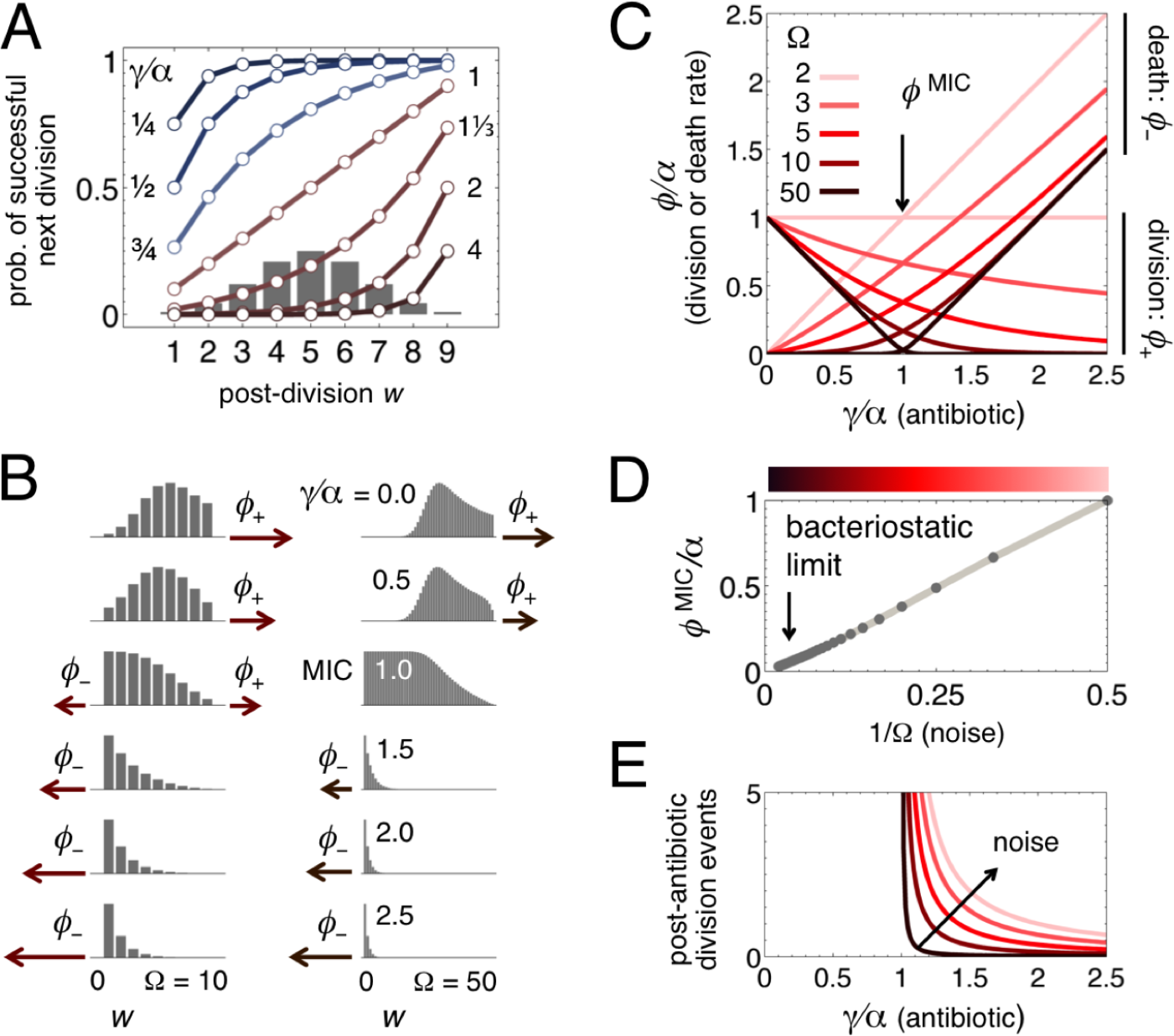
Noise-induced cell division and cell death. (**A**) Two sources of noise: random partitioning and random birth/death of widgets. Immediately after cell division, the number of widgets *w* in a daughter cell follows a binomial distribution (gray histogram). Starting at any widget number, the random birth/death dynamics can take a cell to either boundary. We show the probability that a cell will successfully divide again rather than die (curves; colors represent different values of *γ*/*α* for *Ω* = 10). (**B**) Once sufficient time has passed, distributions of cells over widget number reach a constant shape *f_w_*, as in Eq. 6. We show distributions (gray histograms, scaled to fixed height) as *γ*/*α* = 1 is increased (top to bottom) for two different values of *Ω* (left and right). *γ*/*α* = 1 corresponds to MIC; low *Ω* is high noise, high *Ω* is low noise. The maroon arrows show the resulting rates of cell division (*ϕ*_+_) and cell death (*ϕ*_−_). (**C**) Division rate (*ϕ*_+_; decreasing curves) and death rate (*ϕ*_−_; increasing curves) as a function of antibiotic level (*γ*/*α*) for various values of *Ω*. Darker curves (higher *Ω*) correspond to lower noise. MIC is defined by the point at which *ϕ*_+_ = *ϕ*_−_ = *ϕ*^MIC^. (**D**) As the noise decreases *ϕ*^MIC^ drops, ultimately reaching the “bacteriostatic” limit of zero division and death. 1/*Ω* is the squared coefficient of variation of the binomial distribution in panel (A), and is a convenient measure of noise. (**E**) The total increase in cell number after antibiotic addition, scaled by the original cell number, reflects the number of post-antibiotic cell division events. Below MIC and at MIC itself, the number of division events is infinite; above MIC the population eventually dies out, so the number of division events is finite. Increased noise leads to an increase in the number of division events.

Another way to understand the dynamics is to look at the steady-state distribution of living cells over widget number (Fig. 4B). The more cells there are at the right boundary, the greater the rate of division; the more there are at the left boundary, the greater the rate of death. The addition of antibiotics shifts the entire distribution leftward (Fig. 4B, moving top to bottom). More interestingly, decreasing the noise (increasing *Ω*) narrows the entire distribution away from the boundaries and decreases both division and death rates (compare Fig. 4B, left and right columns).

At the heart of these results is Eq. 7, which relates cell-level parameters to widget-level parameters: it constrains how cell division and death rates (*ϕ*_±_) depend on the antibiotic level (*γ*/*α*). However, this equation on its own does not capture all the details: there are many ways to split *ϕ*_±_ so the constraint of Eq. 7 is satisfied. It is always true that the division rate (*ϕ*_+_) decreases and the death rate (*ϕ*_−_) increases with antibiotic level: at zero antibiotic *γ* = 0 so 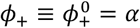, and *ϕ*_−_ = 0; at MIC *α* = *γ* so *ϕ*_+_ = *ϕ*_−_ ≡ *ϕ*^MIC^. But the form of these curves, and in particular the value of *ϕ*^MIC^, depends crucially on the noise level (1/*Ω*). When we plot how *ϕ*_+_ and *ϕ*_−_ depend on *γ*/*α* we see a range of behaviors (Fig. 4C). In the high-noise limit *Ω* = 2 (lightest curves) each cell contains only one widget, and instantaneously divides as soon as this widget replicates. Cell dynamics are thus identical to widget birth/death dynamics, so *ϕ*_+_ = *α, ϕ*_−_ = *γ*, and *ϕ*^MIC^ is high. As the noise is decreased by increasing *Ω* (darker curves), cell division and death curves drop, and *ϕ*^MIC^ also drops. In the low-noise limit of high *Ω* (darkest curves) we approach the bacteriostatic limit: below MIC we see only cell division, above MIC we see only cell death, at MIC we have *ϕ*^MIC^ = 0 so cells neither divide nor die. The classic picture of cell stasis under antibiotic treatment is therefore the low-noise limit of a more general dynamics. Indeed, *ϕ*^MIC^ scales in proportion to the noise level 1/*Ω*. (Fig. 4D), showing that cell division and death at MIC are fundamentally driven by molecular noise.

## Discussion

### Stochastic cell fate choice as a form asymmetric cell division

Suppose we tracked the fate of a single dividing cell in the presence of antibiotics. Due to stochastic effects both its daughters might die or both might divide; the probabilities of these various fates would depend on correlations in inherited components across multiple generations. Our model does not enforce asymmetry in cell division. However, since each widget has an equal chance of being inherited by either daughter cell, these cells can be distinct due to stochastic partitioning. This means the daughters in our model are anti-correlated in their probability of division: the one that inherited more widgets has the higher probability. We could imagine that cells under stress could preferentially partition functional components to one daughter, thus further enhancing survival rates. Such processes have been implicated in the dynamics of bacterial ageing [20: Watve et al., 2006]. Single-cell experiments could be used to track the probability of next division of immediate post-division cells (Fig. 4A). Correlations or anti-correlations in this probability between two daughters would reveal more complex partitioning than we have considered here.

### Continued cell division under antibiotic treatment

At MIC, by definition, for each parent cell on average one of its daughters dies while the other successfully divides. The number of living cells is thus replenished and stays constant, while the number of dead cells continues to increase. For antibiotic levels above MIC, some cells would successfully divide but many more would die, and the population would eventually collapse. (We have not considered persister cells that slow their division under antibiotic treatment [21: Balaban et al., 2004]; these are significant once nearly all cells in the original population have already died). The total increase in OD_600_ post antibiotic treatment measures the number of additional cell divisions before living cells are fully eliminated. Our model predicts that the number of post-antibiotic divisions is significantly amplified by molecular noise (Fig. 4E). This is relevant since each dead cell raises the possibility of sepsis in an infection context. Each cell division is also coupled to a DNA replication event, and represents a chance for new mutations to arise. Molecular noise therefore indirectly increases the possibility of antibiotic resistance.

### Sources of molecular noise

Within our model, the parameter *Ω* plays a nuanced role. Superficially, it measures the number of self-replicating units that trigger cell division. However, it also controls the level of noise in the system. Our model incorporates the intrinsic noise due to the discreteness of widgets, but in practice there will be additional sources of extrinsic noise that increase the rates of division and death [14: Raj & van Oudenaarden, 2008]. In Fig. 1 we showed that the model matches qualitative aspects of our measurements with *Ω* = 10 (note that this is not a numerical fit, just a representative parameter choice). This value of *Ω* predicts *ϕ*^MIC^/*Ω*^0^ = 0.17 (Fig. 4C) whereas the measured value is much higher at *ϕ*^MIC^/*Ω*^0^ 0.6, suggesting that extrinsic noise is significant. To account for this, one could construct more complex models in which the division trigger and noise strength were separately regulated.

### The molecular nature of a widget

Our widget hypothesis captures essential qualitative aspects of the growth and decay of cell populations. However, it does not reveal the nature of a single abstract widget, which is actually a collection of diverse molecules and reactions [15: Jain & Krishna, 2001]. One operational approach to study widgets is via the “Monod curve” which plots cell growth rate as a function of some limiting enzyme concentration. Such a curve often has a threshold enzyme level for non-zero growth, which we can interpret as part of a single self-replicating unit. By comparing the enzyme level at half-maximal growth with this threshold level, we can estimate the number of self-replicating units per cell. Single-cell measurements of *lac* gene expression in *E. coli* suggest this ratio is about 10, to an order of magnitude [11: Kiviet et al., 2014]. We might extend this idea to probe the full molecular composition of cells at the very boundary of death. For each point in composition space, we could operationally define this boundary as the locus of points at which the probability of division drops to some low level, say 1%. The fact that cells can recover from such a state means they must contain at least one self-replicating unit. This suggests a new operational definition of a minimal cell, one that complements existing bottom-up [22: Zhu & Szostak, 2009] and top-down [23: Hutchison et al., 2016] definitions. Monod [24: 1974] would probably have been very comfortable with such an idea.

## Acknowledgements

MT was supported in part by a Wellcome Trust-DBT India Alliance Intermediate Fellowship (500103/Z/09/Z). We thank Sanjay Jain for discussions about bacterial growth laws that stimulated us to start this project.

## Author contributions

MT conceived the project and developed the theoretical model. SC, SD, AS and MT designed the experiments. SV and SC conducted the experiments. SD and MT analyzed the data. MT wrote the manuscript with inputs from all the authors.

## Methods

### Growth and measurement protocols

We grew *Escherichia coli* MG1655 cells, taken from a single colony, overnight in Luria Bertani medium at 37°C. We transferred 50 μL of this culture to 25 mL glucose M9 minimal media in a 100 ml flask at 37°C. The OD_600_ of this culture was monitored until it reached a value of 0.1. At this point, we added kanamycin of the appropriate concentration, and this was defined as the *t* = 0 time point of our measurement. Every 40 minutes, 600 μL of this culture was collected for OD_600_ measurements, and 100 μL was collected for colony counts. We determined both OD_600_ and colony counts using multiple dilutions. Colony counts were measured for four technical replicates; at MIC we performed two biological replicates, each with four technical replicates. Minimal medium composition (100 mL): water, 76.8 mL; 10X M9 salts 10 mL; 20% glucose, 2 mL; 1M CaCl_2_, 10 μL; 100 mM thiamine, 1 mL; 4% casamino acids, 10 mL; 1M MgSO_4_ 200 μL.

### Stochastic model of cell division and death

The transition system shown in Fig. 3D defines a Master Equation:

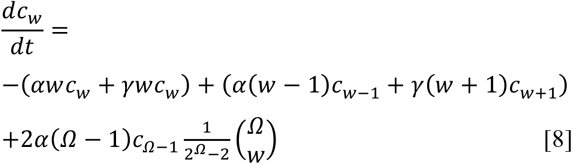

Here, each *c_w_* represents the number of cells (or the normalized probability of cells, depending on the context) with precisely *w* widgets for *w* = 1,…, *Ω* − 1, with the stipulation that *c_Ω_* = 0. The first line corresponds to cells gaining or losing individual widgets. The second line corresponds to the creation of two new daughter cell by the instantaneous division of a cell that hits *Ω* widgets, which happens at rate *α*(*Ω* − 1)*c*_*Ω*−1_. The resulting daughters are defined by *w*′ and *w*″ such that *w*′ + *w*″ = *Ω*. The first factor of 2 accounts for two ways of achieving any given *w*′, in the left or right daughter. The binomial coefficient occurs because each widget has an equal chance of being inherited by either daughter cell. Since a cell divides instantaneously when it hits *Ω* widgets, the usual normalizing factor of 1/2^*Ω*^ is replaced by 1/(2^*Ω*^ − 2): the partitions {*w*′ + *w*″} = {0, *Ω*} or {*Ω*, 0} are ignored since they repeatedly divide until some other partition occurs.

We assume a large number of cells and widgets, so the branching process never goes extinct. If ***c*** = [*c*_1_… *c*_*Ω*-1_]^*T*^ is a column vector, the system of equations Eq. 8 can be written using a transition matrix *A* and solved by matrix exponentiation:

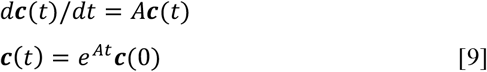

It is convenient to write the vector ***c*** as a product of two components: the number of living cells 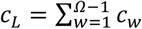, and the normalized distribution *f_w_* of those cells over the different numbers of widgets: ***c***(*t*) ≡ *c_L_*(*t*)***f***(*t*). At long times this distribution approaches the eigenvector of *A* corresponding to its largest eigenvalue: *A**f*** = *λ**f***. Therefore *c_L_*(*t* → *∞*) ∼ *e^λt^*. We can see by direct substitution that *α* - *γ* is an eigenvalue of *A*. Since the number of living cells cannot increase any faster than the number of widgets, we also know this is its largest eigenvalue. Once ***f***(*t*) is determined, Eq. 5 allows us to calculate *ϕ*_+_(*t*) and *ϕ*_−_(*t*) by calculating the rate at which cells cross the right boundary *w* = *Ω* − 1 and the left boundary *w* = 1. By measuring time in units of *α*^−^, we can see that the values *ϕ*_±_/*α* depend only on the ratio *γ*/*α* and on *Ω* (Fig. 4C).

### Probability of division

An immediate post-division daughter cell can have any widget number in the range *w^init^* ∈ {1,…, *Ω* − 1}. Starting from the initial condition 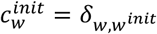, the probability of next division is a first-passage-time problem with absorbing boundaries at *w* = 0 and *w* = *Ω*. This corresponds to a new transition matrix 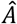 where the binomial partition terms have been removed. We can find 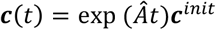 and define

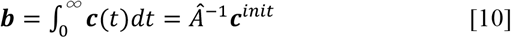

The integrated flux leaving the right and left boundaries are then (Fig. 4A):

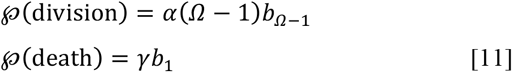
and 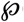 division + 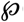 (death) = 1 (Fig. 4A).

